# TET1 interacts directly with NANOG via independent regions containing hydrophobic and aromatic residues

**DOI:** 10.1101/2020.06.29.177584

**Authors:** Raphaël Pantier, Nicholas Mullin, Elisa Hall-Ponsele, Ian Chambers

## Abstract

The DNA demethylase TET1 is highly expressed in embryonic stem cells. Knockout experiments indicate that TET1 is important for lineage commitment, and paradoxically, also for reprogramming to naïve pluripotency. TET1 binds to promoters through a CXXC domain which recognises unmethylated CpG dinucleotides. TET1 also binds to enhancers, presumably via interactions with partner proteins. The transcription factor NANOG interacts with TET1 and is predominantly localised at enhancers in ESCs. Therefore, NANOG may contribute to TET1 biological activity in pluripotent cells. However, the regions of TET1 involved in protein-protein interactions are mostly unknown. Here, we characterise the physical interaction between TET1 and NANOG using embryonic stem cells and bacterial expression systems. TET1 and NANOG interact through multiple binding sites that act independently. Critically, mutating conserved hydrophobic and aromatic residues within TET1 and NANOG abolishes the interaction. Comparative ChIP-seq analysis identifies genomic loci bound by both TET1 and NANOG, that correspond predominantly to pluripotency enhancers. Importantly, around half of NANOG transcriptional target genes are associated with TET1-NANOG co-bound sites. These results indicate a mechanism by which TET1 protein is targeted to specific sites of action at enhancers by direct interaction with a transcription factor.

**Highlights:**

- NANOG and TET1 have regulatory roles in maintaining and reprogramming pluripotency
- TET1 and NANOG interact via multiple independent binding regions
- TET1 and NANOG interactions are mediated by aromatic and hydrophobic residues
- TET1 residues that bind NANOG are highly conserved in mammals
- Co-localisation of TET1 and NANOG on chromatin is enriched at NANOG target genes

## Introduction

Ten-eleven-translocation (TET) family proteins are responsible for active DNA demethylation by oxidation of 5-methylcytosine [1,2] and play important roles during embryonic development and various physiological processes [3]. TET proteins contribute to DNA demethylation in naïve embryonic stem cells (ESCs) [4–7], in particular at enhancers [8–13]. TET protein activity is required both for proper differentiation [14,15] and reprogramming to pluripotency [16–18]. TET1 is the most highly expressed TET family protein both in pluripotent cells and during early development [19–21]. TET1 predominantly binds to promoters via its N-terminal CXXC domain which recognises unmethylated CpG dinucleotides [22–25]. TET1 binding at enhancers in ESCs [26–28], could be mediated by interactions with the pluripotency factors NANOG, PRDM14, OCT4 and SOX2 [29–32]. Interestingly, co-expression of TET1 and NANOG in pre-iPS cells synergistically enhances reprogramming to pluripotency [29]. However, how TET1 might be recruited to chromatin via protein-protein interactions remains poorly understood with little known about the residues involved in protein binding.

Here, the interaction between TET1 and the pluripotency factor NANOG was characterised in ESCs. Co-immunoprecipitations using an array of TET1 truncations and mutants uncovered novel regions involved in protein-protein interactions, both within and outwith the well characterised catalytic domain. Furthermore, alanine mutagenesis identified single residues that show high evolutionary conservation and that contribute to the interaction of TET1 with NANOG. Comparison of TET1 and NANOG ChIP-seq datasets identified genomic loci that are putatively regulated by the TET1-NANOG complex.

## Results

### The TET1 N-terminus interacts directly with NANOG via the evolutionary conserved residues L110 and L114

The TET1 protein expressed in mouse ESCs is composed of 2039 residues. TET1 is characterised by an evolutionary conserved C-terminal catalytic domain, that can be subdivided into a cysteine rich region (residues 1367-1550) and a double stranded beta helix domain (DSBH) (residues 1551-2039) (Figure 1A). TET1 also possesses a CXXC domain (residues 567-608), a DNA binding region [33]. NANOG is a 305 amino acids transcription factor comprising a N-terminal domain (residues 1-95), a DNA binding homeodomain (residues 96-155) and a C-terminal region containing a tryptophan-repeat (WR) (residues 199-243) (Figure 1A). TET1 has been identified as a NANOG-binding protein by independent affinity purification-mass spectrometry analyses [29,30]. To validate the interaction between TET1 and NANOG in pluripotent cells, ESCs were transfected with plasmids expressing (Flag)_3_-TET1 and (HA)_3_-NANOG, respectively. TET1 was immunoprecipitated from nuclear protein extracts using antibodies targeting either a N-terminal (anti-Flag antibody) or a C-terminal (anti-TET1 antibody) epitope (Figure 1B, 1C). Immunoblot analyses showed a substantial enrichment of NANOG following TET1 immunoprecipitation with both antibodies, indicating a physical interaction between these two proteins (Figure 1B, 1C). To determine whether TET1 interacts with NANOG via the TET1 N- or C-terminus, two large (Flag)_3_-tagged TET1 fragments 1-631 and 734-2039 were cloned and expressed in ESCs (Figure 1D), together with NANOG. Following TET1 immunoprecipitation, NANOG was coimmunoprecipitated with both constructs (Figure 1E). As TET1 1-631 and 734-2039 do not contain overlapping residues, these results suggest that TET1 contains at least two NANOG-binding regions that function independently.

**Figure 1.**
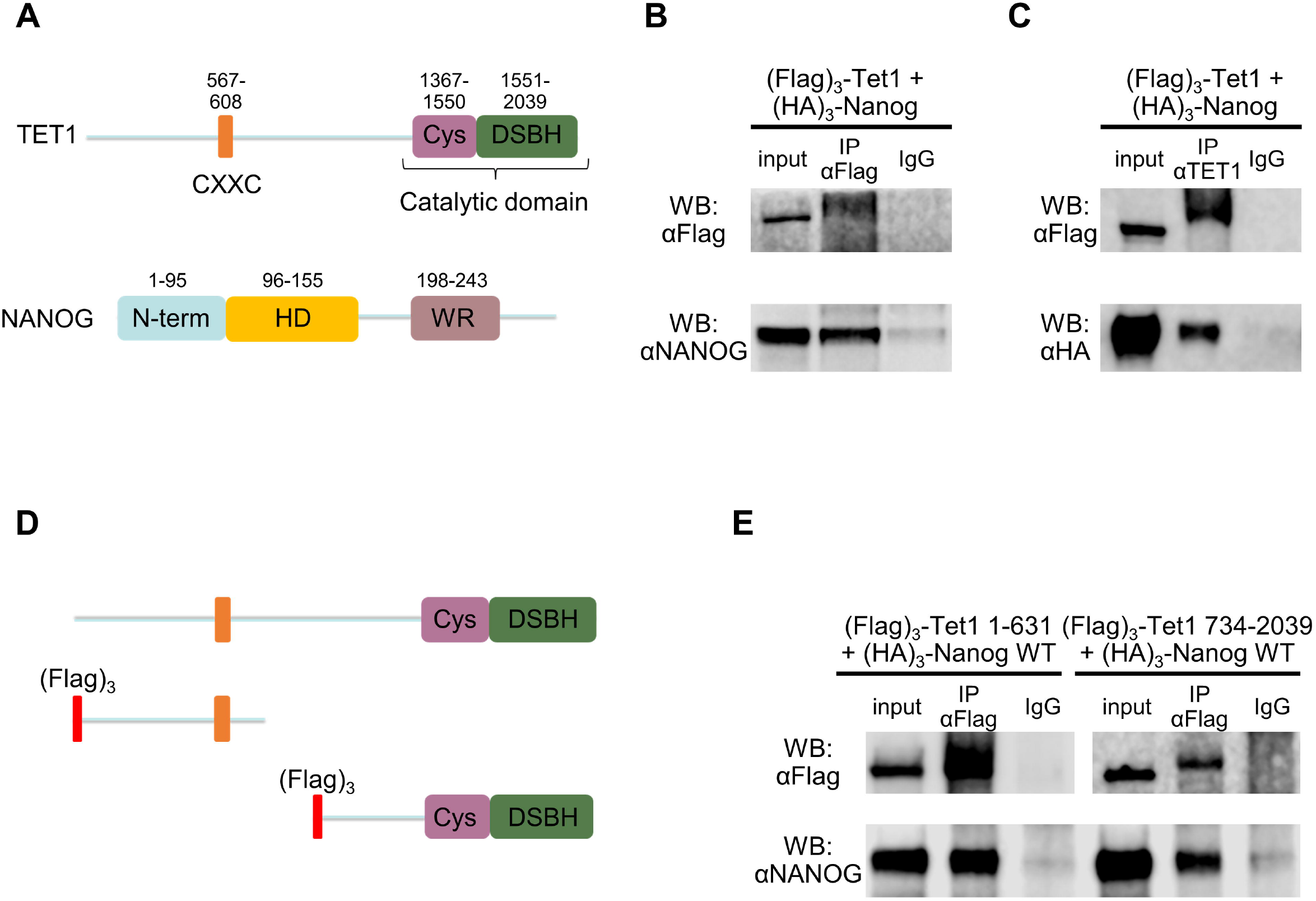
TET1 contains independent NANOG-binding regions. **A.** Diagramatic representation of TET1 and NANOG primary structures; numbers indicate amino acid residues. **B, C.** Co-immunoprecipitation of full-length (Flag)_3_-TET1 (using either anti-Flag (B) or anti-TET1 (C) antibodies) with (HA)_3_-NANOG from E14/T ESCs. Immunoblots were probed with the antibodies indicated on the left. **D, E.** Coimmunoprecipitations of non-overlapping (Flag)_3_-TET1 N- and C-terminal constructs (D) with (HA)_3_-NANOG from E14/T ESCs. Immunoblots (E) were probed with the antibodies indicated on the left.

To begin to explore how NANOG binds to the TET1 N-terminus, (Flag)_3_-tagged TET1 fragments 1-321, 1-215 and 1-108 were cloned and expressed in ESCs (Figure 2A), together with NANOG. NANOG was co-immunoprecipitated with TET1(1-321) and TET1(1-215) but not TET1(1-108) (Figure 2B). To home in on the NANOG-binding site within the TET1 N-terminus and to determine whether the interaction between the TET1 N-terminus and NANOG was direct, both His-tagged NANOG and several MBP-tagged TET1 fragments (1-321, 1-215, 1-165, 1-120, 1-108) were cloned into IPTG-inducible plasmids and expressed in *E.coli* (Figure 2C). MBP-TET1 fragments purified using an amylose resin were examined for co-purifying NANOG by immunoblotting. NANOG co-purified with all TET1 fragments, except TET1(1-108) which showed a dramatically decreased interaction with NANOG (Figure 2D). Importantly, these experiments confirmed a direct physical interaction and narrowed down the first NANOG-binding region to 11 residues (109-120). Protein alignments showed that this region is highly conserved among mammals, indicating a selective pressure for the conservation of these TET1 residues (Figure S1A). However, residues 109-120 of TET1 do not align with TET2 or TET3 proteins (data not shown). To identify which residues are responsible for binding to NANOG, the MBP-TET1 1-120 plasmid construct was modified by alanine substitution of specific amino acids (proline, glutamine, arginine, leucine, serine and valine) within residues 109-120 (Figure 2E). The binding of NANOG to each mutant construct was assessed following bacterial expression and TET1 purification. Strikingly, only the L→A mutant (L110A, L114A) showed a decreased interaction with NANOG, which was reduced to a similar extent as the negative control TET1 1-108 (Figure 2F). Together, these data indicate that one or both of the two evolutionary conserved leucine residues (L110 and/or L114) are in direct physical interaction with NANOG.

**Figure 2.**
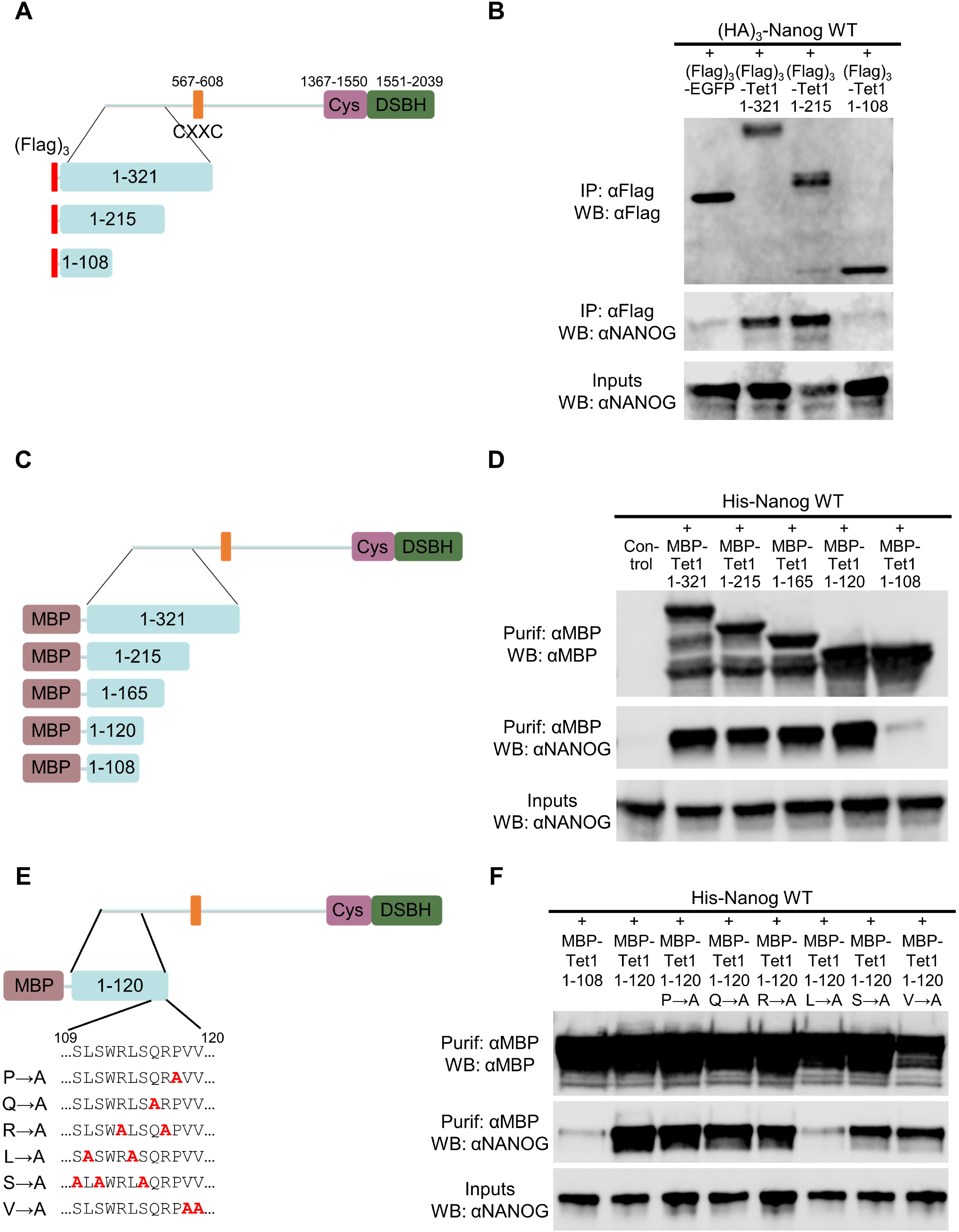
The TET1 N-terminus interacts directly with NANOG via L110 and L114. **A, B.** Co-immunoprecipitations of (Flag)_3_-TET1 N-terminal constructs with (HA)_3_-NANOG from E14/T ESCs. A, fragments of the TET1 N-terminus are shown in the context of full length TET1. B, immunoblots were probed with the antibodies indicated on the left; (Flag)_3_-EGFP was used as a negative control. **C, D.** Co-purification of MBP-tagged TET1 N-terminal constructs with His-NANOG from *E.coli*. C, fragments of the TET1 N-terminus are shown in the context of full length TET1. D, immunoblots were probed with the antibodies indicated on the left. **E, F.** Co-purification of His-NANOG with alanine substitution mutants of MBP-TET1 (1-120) from *E.coli*. E, alanine substitution mutants of TET1 (109-120) are shown in the context of full length TET1. F, immunoblots were probed with the antibodies indicated on the left.

To determine which regions of NANOG interact directly with the TET1 N-terminus, MBP-TET1 1-321 was co-expressed with several His-tagged NANOG truncations in *E.coli* (Figure 3A). Surprisingly, the TET1 N-terminus interacted with three out of four NANOG truncations: 1 160, 91-246, and 194-305 (Figure 3B). However, the TET1 N-terminus showed no interaction with the NANOG homeodomain (Figure 3B). Moreover, while the NANOG WR interacts with MBP-SOX2 (positive control, [30]), MBP-TET1 1-321 showed no physical interaction with the NANOG WR (Figure 3C). Collectively, these results indicate that the TET1 N-terminus interacts with several independent sites on NANOG, and independently of its two most characterised domains: the homeodomain and the WR (Figure 3D).

**Figure 3.**
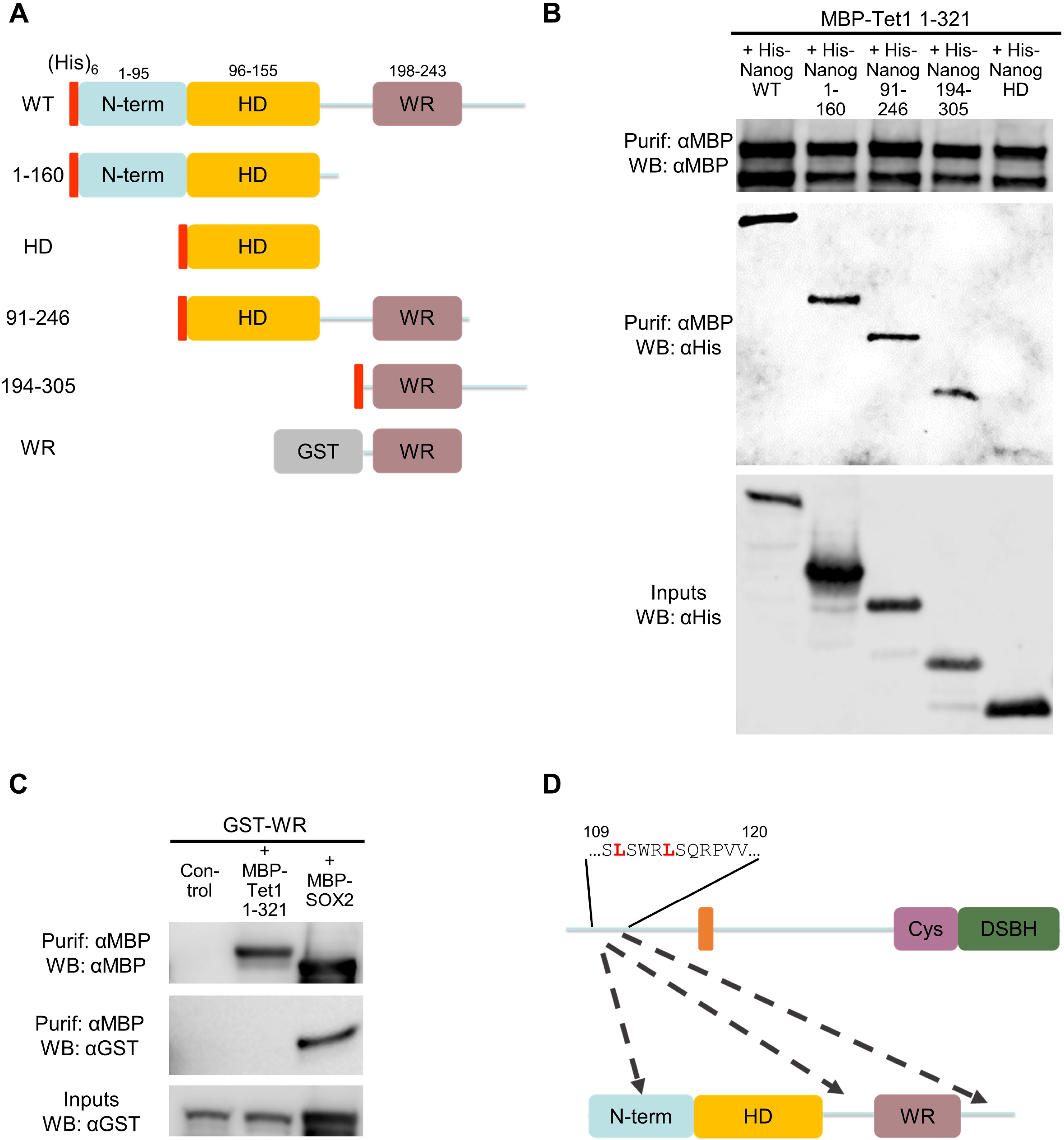
Multiple NANOG regions interact with TET1 N-terminus. **A.** Diagram of the His-tagged (red box) or GST-tagged (grey box) NANOG constructs assayed for binding to MBP-TET1 (1-321) in *E.coli*. **B,** Co-purification of MBP-TET1 1-321 with His-NANOG constructs from *E.coli*. Immunoblots were probed with the antibodies indicated on the left. **C**, analysis of MBP complexes of MBP-TET1 (1-321) or MBP-SOX2 (positive control, [30]) for the presence of GST-WR. Immunoblots were probed with the antibodies indicated on the left. **D.** Diagram of the interactions between the TET1 N-terminus and NANOG, highlighting critical leucines (red). Dashed arrows indicate potential TET1-interacting regions in NANOG.

### The TET1 C-terminus contains two regions that bind NANOG via aromatic interactions

Following the initial observation that TET1 contains >1 independent NANOG-interacting regions (Figure 1E), the TET1 C-terminus was analysed to identify the NANOG-binding residues. Regions of TET1 extending from 734 to varying degrees towards the Cys domain were expressed together with NANOG in ESCs (Figure 4A, S2A). TET1 fragments containing truncations up to residue 1181 (734-1229, 734-1202, 734-1181) were able to bind NANOG, while further C-terminal truncations (734-1155 and 734-1131) abolished interaction with NANOG (Figure S2A, S2B). Analysis of a further truncation (734-1169) narrowed down the second NANOG-binding domain of TET1 to residues 1156-1169 (Figure 4A, 4B). A construct containing this region of TET1 did not interact with NANOG when co-expressed in *E.coli* (Figure S2C and S2D) suggesting that either the interaction observed in ESCs is indirect, or that a direct interaction dependent on post-translational modifications and/or protein folding could not be reproduced in bacteria. However, this interaction does not depend on phosphorylation. While treatment of ESC protein extracts with phosphatase affected the mobilities of both TET1 and NANOG proteins, TET1(734-1169) retained the capacity to bind NANOG (Figure S2E).

**Figure 4.**
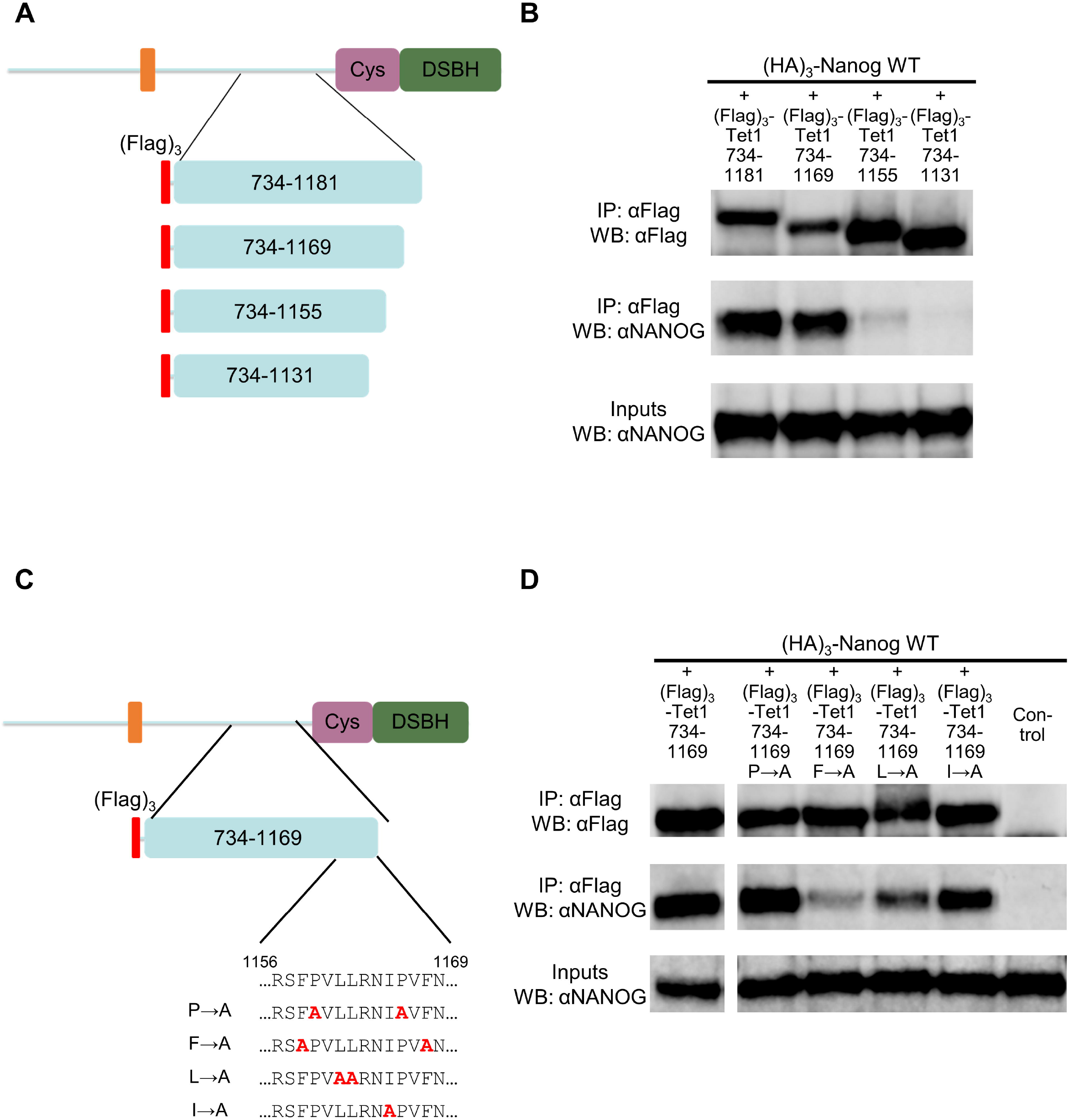
TET1 interacts with NANOG via F1158 and F1168. **A, B.** Co-immunoprecipitation of the (Flag)_3_-TET1 constructs shown in A with (HA)_3_-NANOG from E14/T ESCs. B, immunoblots were probed with the antibodies indicated on the left. **C, D.** Co-immunoprecipitation of alanine substitution mutants of (Flag)_3_-TET1 (734-1169) with (HA)_3_-NANOG from E14/T ESCs. C, alanine substitutions of the amino acids within residues 1156-1169 are shown. D, immunoblots were probed with the antibodies indicated on the left.

Residues 1156-1169 of mouse TET1 have a high similarity to sequences of TET1 proteins from other mammals, with phenylalanine 1158 strictly conserved (Figure S2F). This region did not align with TET2 or TET3 proteins (data not shown). To identify residues that bind NANOG, the expression plasmid encoding TET1 734-1169 was modified by alanine substitution of specific amino acids (proline, phenylalanine, leucine and isoleucine) within residues 1156-1169 (Figure 4C). The F→A mutant (F1158A, F1168A) was the only mutant that showed a decreased interaction with NANOG (Figure 4D). Together, these results indicate that phenylalanine 1158 and/or 1168 are critical for NANOG binding.

The preceding results identified two independent NANOG-binding regions within residues 109-120 (region 1) and 1156-1169 (region 2), respectively in the N- and C-terminal fragments. TET1 fragments containing deletions of these regions did not interact with NANOG compared to the unmutated version (Figure S3A and S3B). Full-length TET1 constructs with deletions in region 1 (Δ1), region 2 (Δ2) or both (Δ1+2) were therefore generated (Figure S3C). A TET1 mutant lacking a low-complexity insert (Δ1733-1901) was used as a control (Figure S3C), as this region has been hypothesised to function in proteinprotein interactions [34]. As expected, NANOG was co-immunoprecipitated with each of the TET1 mutants. Surprisingly however, although the TET1 Δ1+2 double-mutant showed reduced NANOG binding compared to wild-type, binding was not completely eliminated (Figure S3D). This suggests that an additional NANOG-binding region may exist in TET1. To identify this third NANOG-binding region, TET1 expression plasmids were generated combining the double mutation Δ1+2 with increasing C-terminal truncations (Figure 5A). Plasmids with wild-type TET1 coding sequence used as controls allowed assessment of the relative importance of TET1 regions 1 and 2 for NANOG binding. With both wild-type and double-mutant constructs, the TET1-NANOG interaction was dramatically impaired when the TET1 C-terminus was truncated from residue 1547 to 1521 (Figure 5B). Smaller fragments (1-1521, 1-1494, 1-1472 and 1-1379) retained a weak residual interaction with NANOG, which was abolished in double mutants (Δ1 + 2). These results mapped a third NANOG-binding region within TET1 to residues 1522-1547. Most of these residues are strictly conserved in evolution as they are contained within the cysteine-rich catalytic domain (Figure S3E). To identify the residues within this region that bind NANOG, a TET1 construct carrying mutations in regions 1 and 2 (TET1 Δ1 + 2) was further modified by alanine substitution of serine, positively charged or aromatic residues within residues 1522-1547 (Figure 5C). The aromatic→A construct (F1523A, F1525A, W1529A, Y1532A, F1533A, F1538A, F1547A), but not other mutants, reduced the NANOG interaction to a similar extent as the truncated negative control (TET1 1-1521 Δ1+2) (Figure 5D). These data indicate that aromatic residues within TET1 1522-1547 play a critical role for interacting with NANOG in ESCs. Finally, a fulllength TET1 mutant containing mutations in the three NANOG-binding regions identified in this study (Δ109-120 + Δ1132 1202 + 1522-1547 aromatic →A) was generated and tested (Figure 5E). Interestingly, the sequential mutations of regions 1, 2 and 3 within full-length TET1 gradually decreased the interaction with NANOG, to a level comparable to the negative control (Figure 5F, 5G).

**Figure 5.**
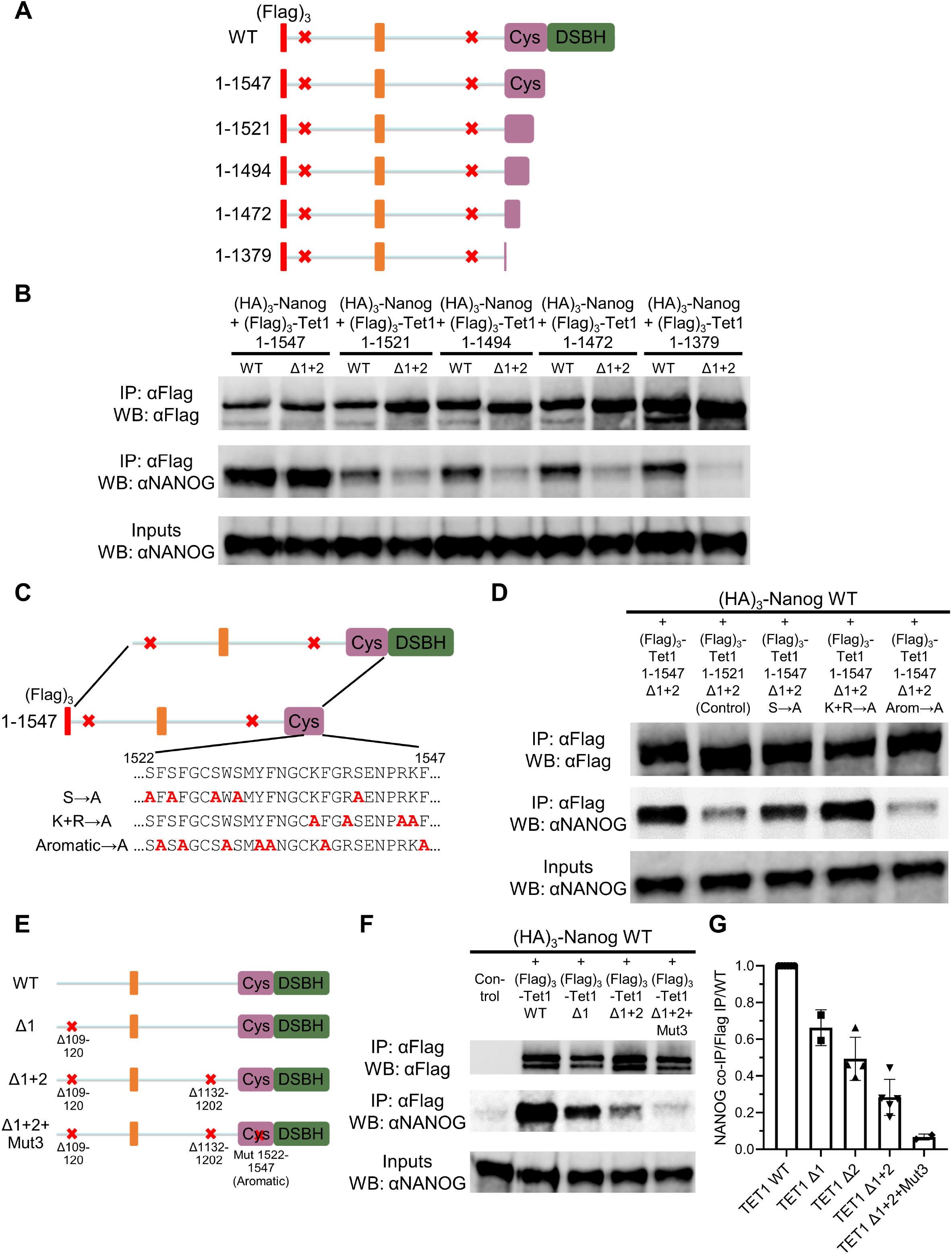
Identification of a third NANOG-binding region and generation of a TET1 triple-mutant unable to interact with NANOG. **A, B.** Co-immunoprecipitation of the indicated (Flag)_3_-TET1 truncations (A) with (HA)_3_-NANOG in E14/T ESCs. TET1 truncations were prepared in parallel in an unmutated (WT) Tet1 plasmid (not shown for simplicity) or one carrying the Δ109-120 + Δ1132-1202 (Δ1+2) mutations (red crosses). B, immunoblots were probed with the antibodies indicated on the left. **C, D.** Co-immunoprecipitations of (Flag)_3_-TET1 mutants (C) with (HA)_3_-NANOG from E14/T ESCs. C, alanine substitution mutants between 1522 and 1547 were prepared in a plasmid expressing TET1(1-1547) with the Δ109-120 + Δ1132 1202 (Δ1+2) mutations (red crosses). D, immunoblots were probed with the antibodies indicated on the left. **E, F, G.** Coimmunoprecipitation of the full-length (Flag)_3_-TET1 mutants indicated (E) with (HA)_3_-NANOG from E14/T ESCs. F, immunoblots were probed with the antibodies indicated on the left. G, quantitation of co-immunoprecipitated NANOG, normalised to TET1 immunoprecipitation levels, relative to wild-type; data points indicate independent experiments and error bars standard deviation.

To identify the NANOG region(s) interacting with TET1 C-terminus, a series of (HA)_3_-tagged NANOG mutants were expressed in ESCs, together with (Flag)_3_-TET1 734-2039 (Figure 6A). Strikingly, only the NANOG mutant lacking the WR region (NANOG ΔWR) showed a reduced interaction with the TET1 C-terminus (Figure 6B). To identify residues within WR responsible for protein-protein interactions, particular amino acids (tryptophan, asparagine, serine and threonine) were substituted by alanine within the WR region of full-length NANOG (Figure 6C). Only the W→A mutant showed a decreased interaction with the TET1 C-terminus (Figure 6D), demonstrating a key role for tryptophans in the interaction with Tet1 C-terminus. Together, these experiments demonstrate that NANOG interacts with the TET1 C-terminus via aromatic residues conserved in both proteins (Figure 6E).

**Figure 6.**
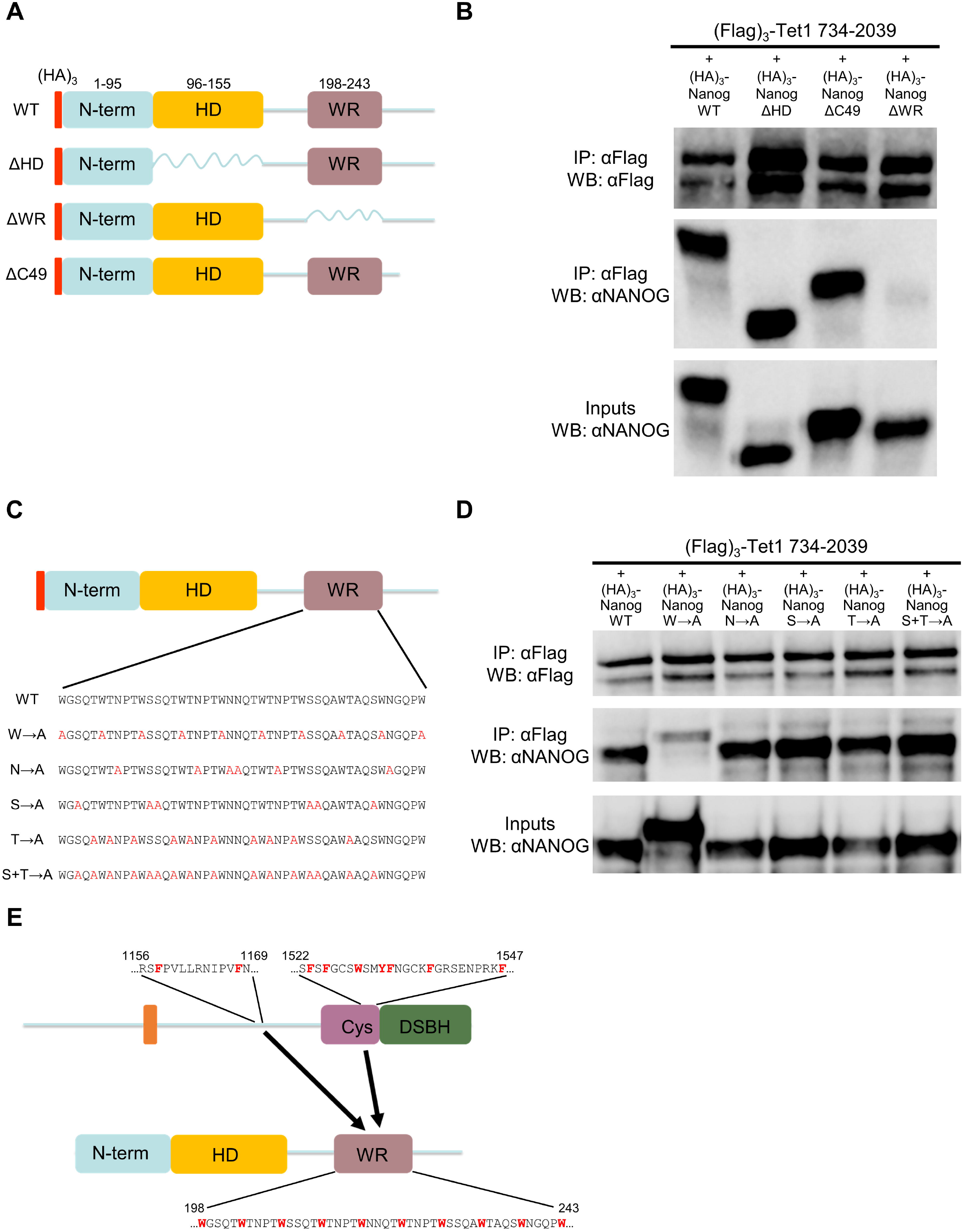
Tryptophan residues within NANOG WR interact with TET1 C-terminus. **A, B.** Co-immunoprecipitations of (Flag)_3_-TET1 (734-2039) with the indicated (HA)_3_-NANOG deletion mutants (A) from E14/T ESCs. B, immunoblots were probed with the antibodies indicated on the left. **C, D.** Co-immunoprecipitation of (Flag)_3_-TET1 (734-2039) with (HA)_3_-NANOG or the derivative alanine substitution mutants within the WR from E14/T ESCs. Only amino acids within NANOG WR region were mutated to alanine (C). Immunoblots (D) were probed with the antibodies indicated on the left. **E.** Diagram of the interactions between the TET1 C-terminal domains and NANOG, highlighting residues critical for the interaction (red).

### TET1 and NANOG co-bind a subset of pluripotency enhancers associated with NANOG transcriptional target genes

Although TET1 and NANOG interact directly in ESCs, the relationship between the two proteins on chromatin remains unclear. To identify genomic sites potentially regulated by the TET1-NANOG complex, published TET1 and NANOG ChIP-seq datasets were compared. TET1 ChIP-seq peaks from two independent datasets [23,24] showed an overlap of 13,279 “high confidence” TET1 binding sites (Figure S4A). A similar analysis of two NANOG ChIP-seq studies [35,36] identified 24,357 “high confidence” NANOG ChIP-seq peaks (Figure S4B). Subsequently, TET1 and NANOG ChIP-seq signals were visualised at high confidence NANOG and TET1 binding sites, respectively. Interestingly, TET1 is enriched at the centre of a large proportion of NANOG binding sites in ESCs, and this signal is abolished upon Tet1 knockdown (Figure 7A). In contrast, NANOG is enriched only at a small proportion of TET1 binding sites in ESCs (Figure S4C). Consistent with this low level of co-enrichment, the stringent intersection of “high confidence” TET1 and NANOG ChIP-seq peaks identified only 2,003 sites bound by both TET1 and NANOG (Figure S4D). As a first inspection, TET1-NANOG peaks were crossed with relevant genomic features, showing a large proportion of sites corresponding to ESC enhancers [37] (65%) and a smaller proportion overlapping with CpG islands [38] (22%) (Figure 7B). Remarkably, *de novo* motif analysis identified the SOX2/OCT4 composite motif at TET1-NANOG co-bound sites (Figure 7C). Following these observations, further analyses were performed to characterise genes associated with TET1-NANOG ChIP-seq peaks. Gene ontology analysis identified groups of genes associated with pluripotency among the top categories, such as “stem cell population maintenance”, “cellular response to leukemia inhibitory factor” and “cell fate specification” (Figure S4E). Importantly, TET1-NANOG ChIP-seq peaks were found within or in proximity to 48% of NANOG transcriptional target genes [39] (Figure 7D and Table 1). Visual inspection of these loci showed enrichment of TET1 and NANOG ChIP-seq signals at known enhancers and putative cis-regulatory elements (Figure S4F). Together, these results suggest that the TET1-NANOG complex regulates a significant subset of NANOG target genes.

**Figure 7.**
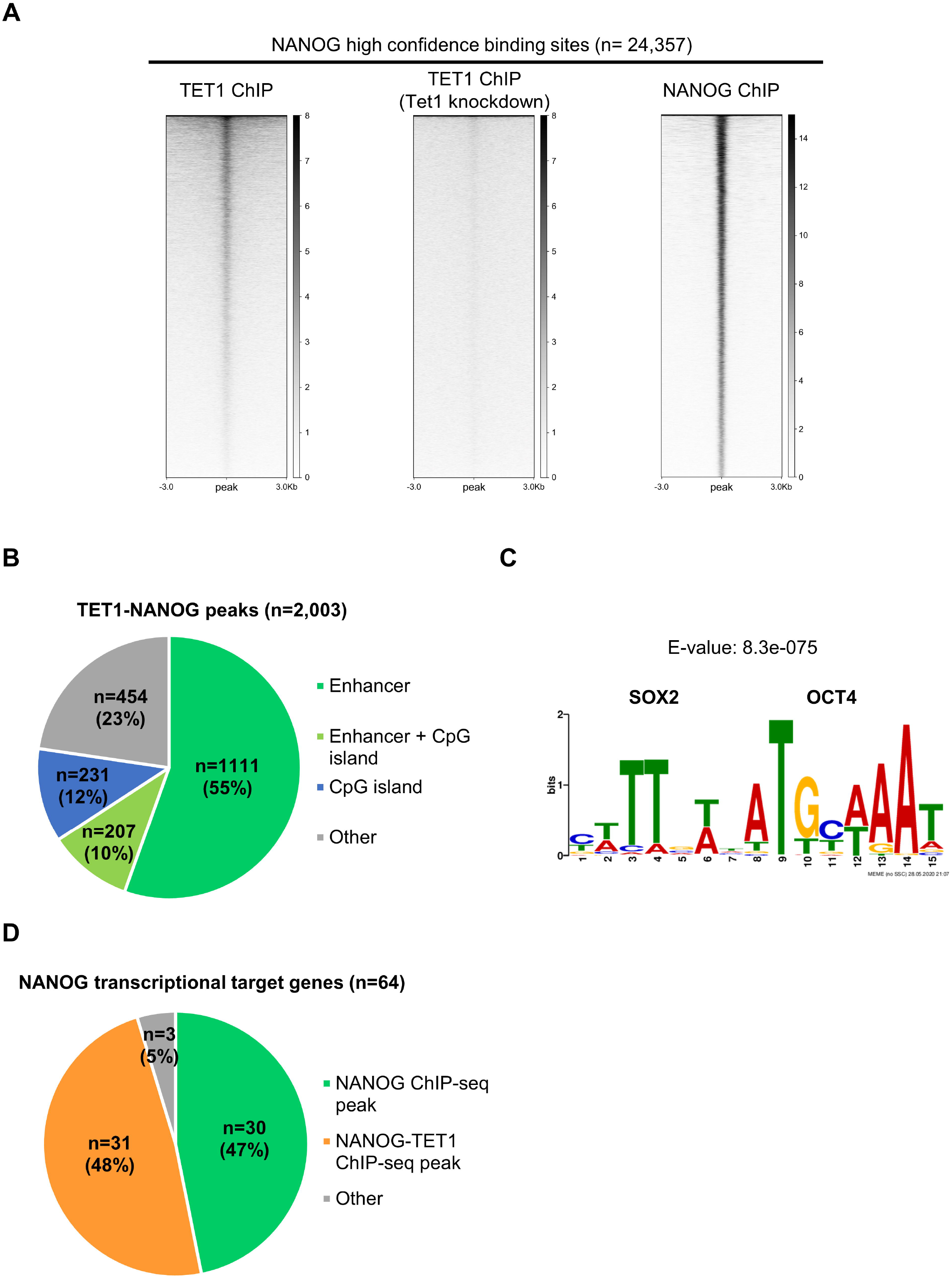
Identification of TET1-NANOG co-bound sites on chromatin in ESCs. **A.** TET1 and NANOG ChIP-seq signals at NANOG “high confidence” binding sites, as defined in Figure S4B. TET1 ChIP-seq in ESCs treated with Tet1 shRNA (knockdown) was used as a negative control. **B.** Pie chart showing the portion of TET1-NANOG co-bound sites (see Figure S4D) overlapping with ESC enhancers and CpG islands. **C.** *De novo* motif analysis performed on TET1-NANOG co-bound sites, showing the most significant binding motif and its respective E-value. **D.** Pie chart showing the portion of NANOG transcriptional target genes with NANOG or TET1-NANOG co-bound sites.

**Table 1.** NANOG transcriptional target genes associated with TET1-NANOG co-bound sites

| ***Activated genes*** | ***Repressed genes*** |
| --- | --- |
| *B4gat1* | *Ctbp2* |
| *Cdc42ep4* | *Dusp1* |
| *Cpsf4l* | *Edn2* |
| *En1* | *Fzd7* |
| *Esrrb* | *Hmces* |
| *Igf2bp2* | *Lefty2* |
| *Kit* | *Lpar6* |
| *Klf4* | *Nid2* |
| *Lmo4* | *Otx2* |
| *Manba* | *Raet1e* |
| *Mras* | *Rbm47* |
| *Plekhg3* | *Rbpms* |
| *Plpp1* | *Smagp* |
| *Setd1b* | *Socs3* |
| *Sp5* | *Tcf15* |
| *Tmem51* |  |

## Discussion

TET1 [19] and NANOG [40,41] are both expressed in the inner cell mass of the blastocyst, which is modelled *in vitro* by ESCs. TET1 and NANOG are also co-expressed in the postimplantation epiblast [20,42,43] and in developing primordial germ cells [43,44]. Loss of either TET1 or NANOG compromises germline development [45–48]. Therefore, the TET1-NANOG interaction reported here may function not only at pre-implantation stages but also during later development.

Alanine substitution mutagenesis identified aromatic and hydrophobic residues that mediate the interaction between TET1 and NANOG. In NANOG, tryptophans within the WR are critical for the biological function of NANOG and mediate NANOG homodimerization and binding to SOX2 aromatic residues [49,50,30,51]. Tryptophan residues within the NANOG WR also interact with aromatic residues in the TET1 C-terminus, suggesting an interaction by aromatic stacking. The present work also demonstrates a direct, WR-independent interaction between NANOG and the TET1 N-terminus, indicative of novel protein interaction sites in NANOG.

TET1 has previously been reported to interact with the SIN3A PAH1 domain by amphipathic helix formation [52] and with OGT via C-terminal TET1 residues [53]. Other TET1-interacting proteins have been identified [29,31,32,54–57]. These include thymine DNA glycosylase, which binds TET1 through at least two sites [54]. However, apart from SIN3A and OGT, the residues mediating these interactions have not been defined. The present work demonstrates a direct physical interaction between TET1 and NANOG. Strikingly, the TET1-NANOG interaction involves multiple independent binding regions between both proteins. The three NANOG interaction domains on TET1 (residues 109-120, 1156-1169 and 1522-1547) have not been characterised in other protein interaction studies. This adds new information about TET1 function and suggests that TET1 could also interact with other proteins through multiple binding sites. Interestingly, one of the NANOG-interaction domains (1522-1547) includes residues that interact with DNA, that lie adjacent to the TET1 catalytic domain and that are conserved in TET2 and TET3 [58]. We have recently identified two binding regions in TET2 that interact with NANOG and one of these includes residues homologous to TET1 1522-1547 [21]. Importantly, this region (1522-1547) also contains residues that bind methylated CpG [58]. The TET1-NANOG interaction seems to be DNA-independent since the interaction is seen in DNase-treated protein extracts, and since the interaction is unaffected by deletion of the NANOG homeodomain or the TET1 CXXC domain. Therefore, binding of NANOG to TET1 residues 1522-1547 could modulate the interaction of the catalytic domain of TET1 with DNA that depends on these residues. It will therefore be of interest to determine whether the interaction with NANOG modulates TET1 catalytic activity.

Here, comparative analysis of TET1 and NANOG ChIP-seq datasets identified a subset of genomic loci co-bound by TET1 and NANOG in ESCs. Interestingly, the majority of these sites correspond to pluripotency enhancers and are enriched for the SOX2/OCT4 motif [59]. About half of NANOG target genes have a TET1-NANOG peak nearby, suggesting that TET1 may act cooperatively with NANOG to regulate transcription [60]. NANOG target genes that have an associated TET1-NANOG peak include genes that are either activated or repressed by NANOG in ESCs. Potentially, TET1 could modulate transcription by demethylating enhancer DNA [18,29,61]. Furthermore, TET1 may regulate the expression of NANOG target genes by recruiting the SIN3A co-repressor complex at these loci [23,62,63]. However, further investigation will be required to unravel the mechanisms by which enhancers may be co-regulated by TET1 and NANOG and to distinguish action at positively and negatively regulated NANOG target genes.

## Materials and methods

### Molecular cloning

TET1 open reading frame was subcloned into pPYCAG plasmids for exogenous expression of Flag-tagged proteins in embryonic stem cells [40]. TET1 open reading frame was subcloned into pRSFDuet plasmids (Novagen) for exogenous expression of MBP-tagged proteins in *E. coli*. TET1 truncations and mutants were obtained by cloning PCR products or synthetic DNA fragments (Integrated DNA Technologies, Inc.) using Gibson Assembly [64].

### Cell culture

E14/T mouse embryonic stem cells were used in this study, as they constitutively express the polyoma large T antigen and can therefore propagate pPYCAG plasmids carrying the polyoma origin of replication [40]. ESCs were cultured in a 37°C/7% CO_2_ incubator on gelatin-coated plates. Composition of the culture medium: Glasgow minimum essential medium (Sigma-Aldrich, cat. G5154), 10% fetal bovine serum, 1× L-glutamine (Thermo Fisher Scientific, cat. 25030024), 1x sodium pyruvate (Thermo Fisher Scientific, cat. 11360039), 1× MEM non-essential amino acids (Thermo Fisher Scientific, cat. 11140035), 0.1mM 2-Mercaptoethanol (Thermo Fisher Scientific, cat. 31350010), 100U/ml LIF (made inhouse).

To overexpress tagged proteins for co-immunoprecipitations, 3×10^6^ E14/T ESCs were transfected with pPYCAG plasmids of interest using Lipofectamine 3000 (Thermo Fisher Scientific, cat. L3000008). Transfections were performed in 10cm dishes following manufacturer’s instructions. E14/T ESCs were harvested 24h after transfection for protein extract preparation.

### Preparation of nuclear protein extracts from embryonic stem cells

ESCs were washed twice with PBS, trypsinised and pelleted (5 min, 400g, 4°C) before lysis in swelling buffer (5 mM PIPES pH8.0, 85mM KCl) freshly supplemented with 1x protease inhibitor cocktail (Roche, cat. 04693116001) and 0.5% NP-40. After 20 min on ice with occasional shaking, nuclei were pelleted (10 min, 500g, 4°C) and resuspended in 1 ml of lysis buffer (20mM HEPES pH7.6, 350mM KCl, 0.2mM EDTA, 1.5mM MgCl_2_, 20% glycerol) freshly supplemented with 0.2% NP-40, 0.5 mM DTT, and 1x protease inhibitor cocktail. The material was transferred into no-stick microtubes (Alpha Laboratories, cat. LW2410AS) and supplemented with 150 U/ml of Benzonase nuclease (Millipore, cat. 71206). Samples were incubated on a rotating wheel for 30 min at 4°C and centrifuged (16,000g, 30 min, 4°C) to remove any precipitate. Nuclear proteins extracts were stored at −80°C, or used directly for immunoprecipitation or immunoblot. 30-50μl of protein extract was used as input material and boiled in Laemmli buffer for 5min at 95°C.

### Immunopurification of Flag-tagged proteins from nuclear protein extracts

To immunoprecipitate TET1, 5μg of anti-Flag (Sigma-Aldrich, cat. F3165) or anti-TET1 (Millipore, cat. 09-872) antibody was added to protein extracts. For negative controls, 5μg of normal IgG (Santa Cruz) were added to protein extracts. Samples were incubated overnight at 4°C on a rotating wheel. 30μl of beads coupled with ProteinA or ProteinG (GE Healthcare 4 Fast Flow Sepharose), previously blocked with 0.5mg/ml chicken egg albumin (Sigma-Aldrich), were added to each sample, followed by a 2h incubation at 4°C on a rotating wheel. Beads were washed 5 times with lysis buffer (20mM HEPES pH7.6, 350mM KCl, 0.2mM EDTA, 1.5mM MgCl_2_, 20% glycerol) freshly supplemented with 0.5% NP-40 and 0.5 mM DTT. Between each wash, samples were centrifuged at 4°C for 1min at 2,000rpm. After the final wash, beads were resuspended in Laemmli buffer and boiled for 5min at 95°C

As an alternative strategy to immunoprecipitate Flag-tagged proteins, 30μl of anti-Flag magnetic beads (Sigma-Aldrich, cat. M8823) was added to each protein extract. Samples were incubated on a rotating wheel for 2h at room temperature. Following three washes with PBS using a magnet (Thermo Fisher Scientific, cat. 12321D), magnetic beads were resuspended in Laemmli buffer and boiled for 5min at 95°C. Samples were stored at −20°C or analysed directly by immunoblot.

### Preparation of protein extracts from bacterial pellets

Chemically competent BL21(DE3) *E.coli* (NEB, cat. C2527I) were transformed with pRSF bacterial expression plasmids of interest. A single colony was inoculated in LB medium supplemented with appropriate antibiotics and incubated overnight in a 37°C shaker (225rpm). The overnight culture was diluted (1/50) in a 50ml flask containing 50ml of LB medium supplemented with appropriate antibiotics and incubated in a 37°C shaker (225rpm) until the culture reached the exponential phase (≈3h, OD_600_:0.5–0.7). 1mM IPTG was added to the culture to initiate protein expression, and cells were transferred in an 18°C shaker (225rpm) for 6h. Bacterial pellets were collected by centrifugation (5,000g, 10min) and stored at −20°C until protein extraction.

To prepare protein extracts, bacterial pellets were resuspended in 5ml of cold protein extraction buffer (25mM Tris-HCl pH 8.0, 200mM NaCl), and sonicated 3×1min on ice. Samples were centrifuged (16,000g, 30 min, 4°C) to remove insoluble material. Bacterial protein extracts were stored at 4°C or used directly for protein purification. 30-50μl of protein extract was used as input material and boiled in Laemmli buffer for 5min at 95°C.

### Purification of MBP-tagged proteins from bacterial extracts

To purify MBP-tagged proteins, each bacterial protein extract was loaded into a gravity flow column containing 600μl of amylose resin. The resin was washed once with cold protein extraction buffer (25mM Tris-HCl pH 8.0, 200mM NaCl) and MBP-tagged proteins were eluted in 500μl of cold protein extraction buffer (25mM Tris-HCl pH 8.0, 200mM NaCl) supplemented with 10mM Maltose. 50μl of eluate was boiled in Laemmli buffer for 5min at 95°C.

### Immunoblot

Protein samples were loaded into Bolt 10% Bis-Tris Plus Gels (Thermo Fisher Scientific, cat. NW00102BOX) with 1x Bolt MOPS SDS running buffer (Thermo Fisher Scientific, cat. B0001). 10μl of SeeBlue Plus2 pre-stained protein standard (Thermo Fisher Scientific, cat. LC5925) was used to visualize protein molecular weight. The electrophoresis was performed at 160V for 1h. Proteins were transferred overnight at 4°C onto a nitrocellulose membrane (150mA constant current) with transfer buffer (25mM Tris, 0.21M glycine, 10% methanol). The membrane was blocked for 1h at room temperature with 10% (w/v) non-fat skimmed milk dissolved in PBS supplemented with 0.1% Tween. Then, the membrane was incubated for 1h at room temperature with primary antibodies diluted to the working concentration in 5% (w/v) non-fat skimmed milk dissolved in PBS supplemented with 0.1% Tween. The membrane was washed 3 times with PBS supplemented with 0.1% Tween, and incubated for 2h at room temperature with LI-COR IRDye conjugated secondary antibodies diluted 1:5,000 in 5% non-fat skimmed milk dissolved in PBS supplemented with 0.1% Tween. The membrane was finally washed 3 times with PBS supplemented with 0.1% Tween and analysed with the LI-COR Odyssey FC imaging system.

### Antibodies

| Antibody | Reference | Working dilution (immunoblot) |
| --- | --- | --- |
| Flag | Sigma-Aldrich, cat. F3165 | 1:5,000 |
| HA | Covance, cat. MMS-101P | 1:5,000 |
| Nanog | Bethyl, cat. A300-397A | 1:2,000 |
| GST | Abcam, cat. ab92 | 1:2,000 |
| His tag | Abcam, cat. ab18184 | 1:2,000 |
| MBP | NEB, cat. E8032S | 1:10,000 |

### Protein alignments

To identify evolutionary conserved residues, TET1 protein sequences from various mammalian and non-mammalian species were downloaded from UNIPROT (https://www.uniprot.org/) and aligned using ESPript (http://espript.ibcp.fr) [65].

### ChIP-seq analysis

ChIP-seq datasets were analysed using the Galaxy platform (https://usegalaxy.org) [66]. Details concerning the bioinformatic workflow are available at the following address: https://usegalaxy.org/u/raf4579/w/workflow-chip-seq-1. Raw sequencing data (FASTQ files) was downloaded from publicly available databases NCBI’s Gene Expression Omnibus or ArrayExpress. Quality control was performed using the software “FastQC” (Babraham Bioinformatics). Samples were filtered to remove contaminating adapter sequences and low-quality reads (cut-off quality score >20.0). Reads were mapped to the mouse mm9 reference genome using “Bowtie2” (BAM file output) [67]. Reads were mapped only to a unique genomic location (k=1). ChIP-seq peaks were called using the software “MACS2” (BED file output) [68]. The immunoprecipitated sample was compared to the genomic input for identifying statistically significant binding sites (qvalue 0.05). If replicates were available, only ChIP-seq peaks shared between replicates were considered for further analyses. For the analysis of NANOG ChIP-seq datasets, the algorithm optimised for “narrow peaks” was used. For the analysis of TET1 ChIP-seq datasets, the algorithm optimised for “broad peaks” was used. To visualise ChIP-seq datasets on a genome browser, mapped reads (BAM files) were converted into bigWig files using “Deeptools” [69]. Data was normalised in “Reads Per Kilobase Million” (RPKM) to allow the comparison between ChIP-seq datasets. Genomic snapshots were taken using the genome viewer “IGV” [70]. To visualise ChIP-seq datasets as heatmaps, the software “Deeptools” was used [69]. To perform *de novo* motif analysis on ChIP-seq datasets, the DNA sequences corresponding to each ChIP-seq peak were extracted (FASTA file output) and analysed using the “MEME” software [71]. Motifs between 5 and 25bp, enriched with a E value <0.05, were identified. These results were compared to known protein motifs in the JASPAR database [72]. Assignment of genes to ChIP-seq peaks and gene ontology analysis were performed using the “Genomic Regions Enrichment of Annotations Tool” (GREAT).

## Accession numbers

UniProt Knowledgebase (UniProtKB) accession numbers: **<u>E9Q9Y4</u>** (Mouse TET1 protein sequence), **<u>Q80Z64</u>** (Mouse NANOG protein sequence).

NCBI Gene Expression Omnibus accession numbers: **<u>GSE24841</u>** (TET1 ChIP-seq), **<u>GSE26832</u>** (TET1 ChIP-seq), **<u>GSE44286</u>** (NANOG ChIP-seq).

ArrayExpress accession numbers: **<u>E-MTAB-1617</u>** (NANOG ChIP-seq).

## CRediT authorship contribution statement

**Raphaël Pantier:** Conceptualization, Methodology, Investigation, Writing - Original Draft, Writing - Review & Editing Visualization. **Nicholas Mullin:** Conceptualization, Methodology, Supervision, Writing - Review & Editing. **Elisa Hall-Ponsele:** Investigation. **Ian Chambers:** Conceptualization, Writing - Original Draft, Writing - Review & Editing, Supervision, Project administration, Funding acquisition

## Acknowledgements

We thank Elisa Barbieri for constructive comments on the manuscript. We are grateful to Kristian Helin (University of Copenhagen and Memorial Sloan Kettering Cancer Center) for sharing Tet1 expression plasmids. This work was funded by a UK Medical Research Council Grant MR/L018497/1 to IC. RP was supported by a UK Medical Research Council PhD Fellowship.

## Abbreviations

DSBH: double stranded beta helix domain
ESC: embryonic stem cell
OGT: O-linked N-acetylglucosamine transferase
TET: Ten-eleven-translocation
WR: tryptophan-repeat

## Supplementary figure captions

**Supplementary Figure 1.**
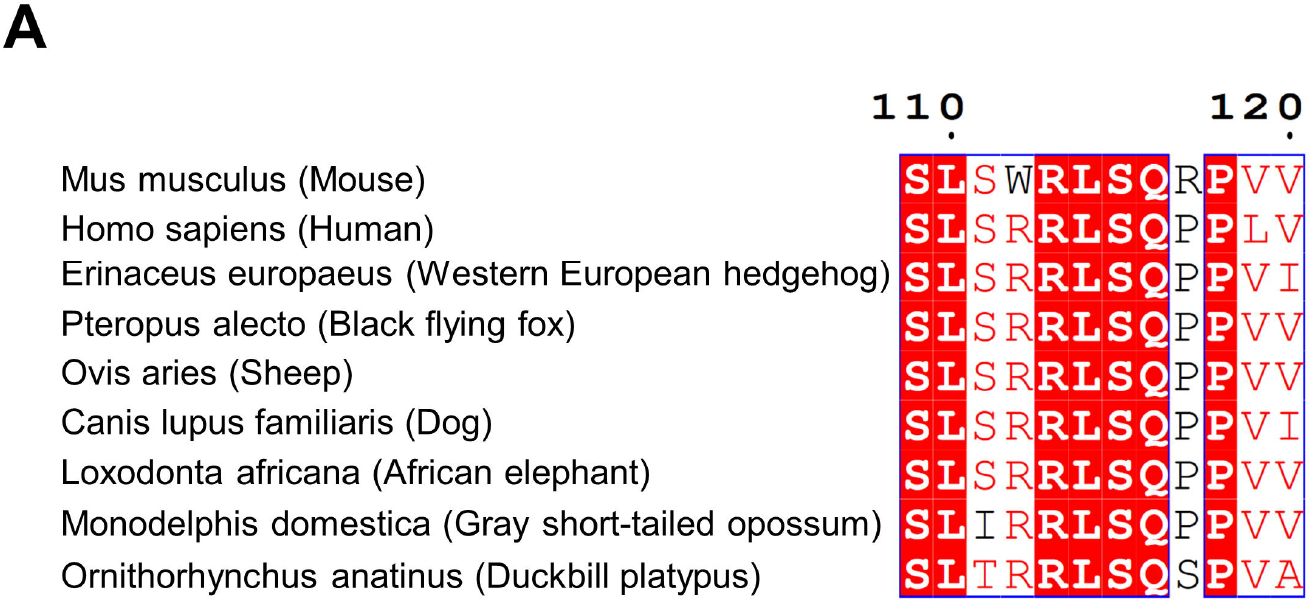
(related to Figure 2) **A.** Alignment of the NANOG-interacting domain I in mammalian TET1 proteins, centred on mouse TET1 residues (109-120). Identical residues are white on a red background; conservative substitutions found in several mammalian species are in red.

**Supplementary Figure 2.**
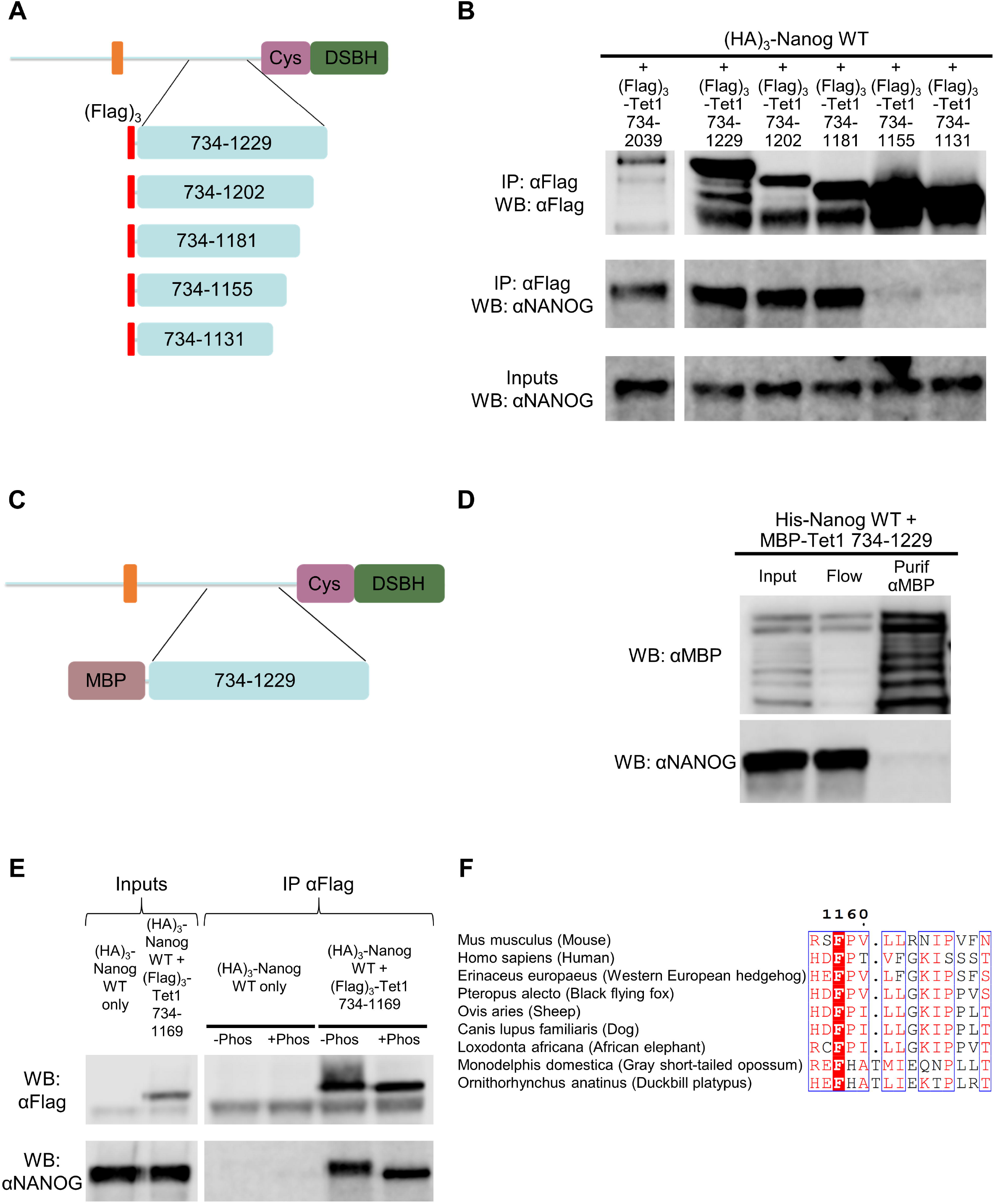
(related to Figure 4) **A, B.** Co-immunoprecipitation of C-terminal truncations of (Flag)_3_-TET1(734-1229) (A) with (HA)_3_-NANOG from E14/T ESCs. B, immunoblots were probed with the antibodies indicated on the left. **C, D.** Co-purification of MBP-TET1(734-1229) (C) with His-NANOG from *E.coli*. D, immunoblots were probed with the antibodies indicated on the left. **E.** Coimmunoprecipitations of (Flag)_3_-TET1(734-1169) with (HA)_3_-NANOG from E14/T ESCs with (+Phos) or without (-Phos) phosphatase treatment. Immunoblots were probed with the antibodies indicated on the left. **F.** Alignment of NANOG-interacting domain II of TET1 proteins from the indicated mammals, centred on mouse TET1 residues (1156-1169). Identical residues are white on a red background; conservative substitutions found in several mammalian species are in red.

**Supplementary Figure 3.**
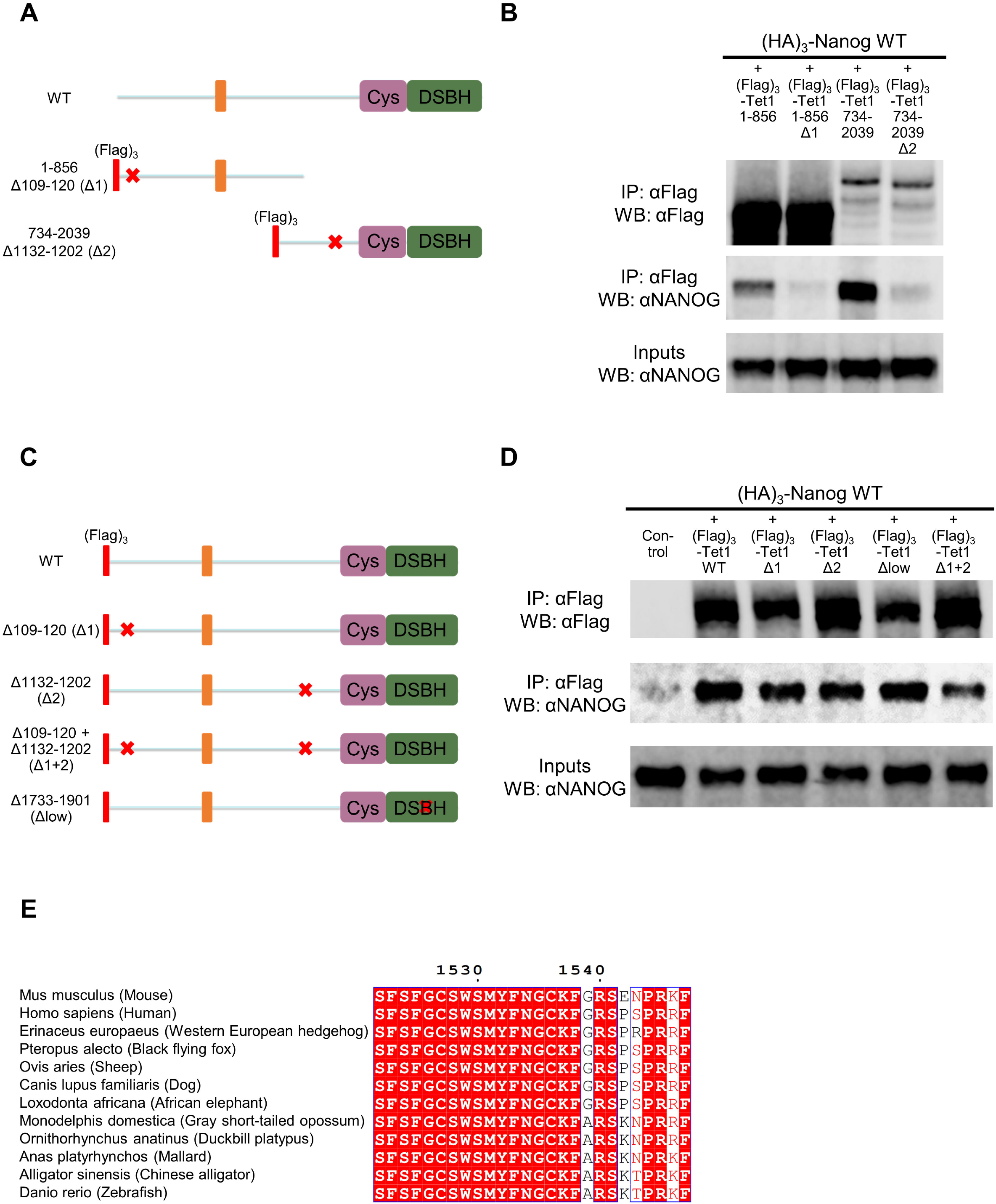
(related to Figure 5) **A, B.** Co-immunoprecipitation of (Flag)_3_-TET1 truncations (A) with (HA)_3_-NANOG from E14/T ESCs. TET1 truncations were prepared in plasmids carrying the Δ109-120 (Δ1) or Δ1132-1202 (Δ2) mutations indicated (red crosses). B, immunoblots were probed with the antibodies indicated on the left. **C, D.** Co-immunoprecipitation of full-length (Flag)_3_-TET1 mutants (C) with (HA)_3_-NANOG in E14/T ESCs. TET1 constructs carried the Δ109-120 (Δ1), Δ1132-1202 (Δ2) or Δ1733-1901 (Δlow) mutations indicated (red crosses). D, immunoblots were probed with the antibodies indicated on the left. **E.** Alignment of the NANOG-interacting domain III of TET1 from the indicated species, centred on mouse TET1 residues (1522-1547). Identical residues are white on a red background; conservative substitutions found in several mammalian species are in red.

**Supplementary Figure 4.**
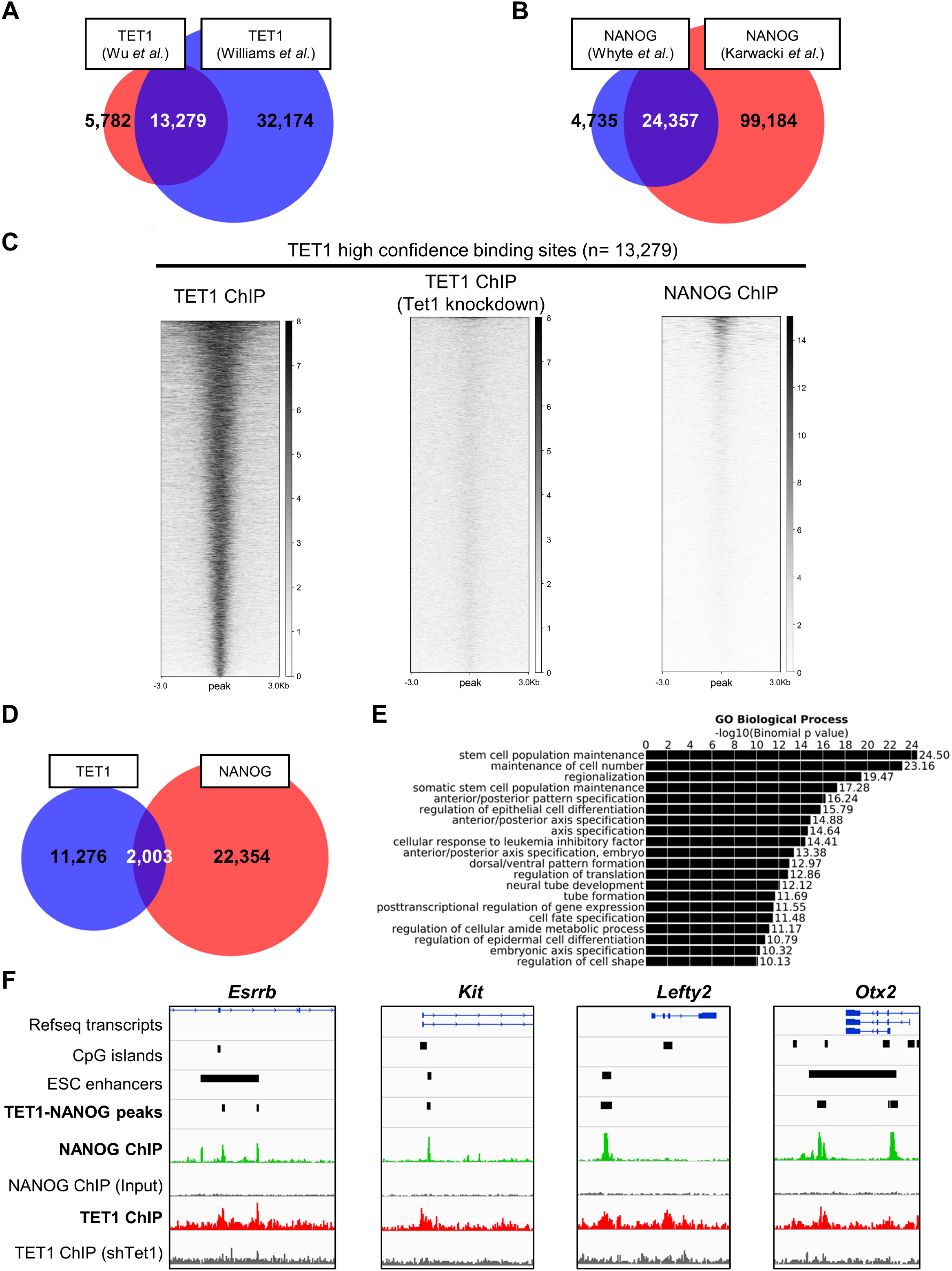
(related to Figure 7) **A.** Venn diagram showing the overlap of TET1 ChIP-seq peaks between two published datasets in mouse ESCs [23,24]. High confidence TET1 binding sites are shared between datasets (n= 13,279). **B.** Venn diagram showing the overlap of NANOG ChIP-seq peaks between two published datasets in mouse ESCs [35,36]. High confidence NANOG binding sites are shared between datasets (n= 24,357). **C.** TET1 and NANOG ChIP-seq signal at TET1 “high confidence” binding sites, as defined in Figure S4A. TET1 ChIP-seq in ESCs treated with Tet1 shRNA (knockdown) was used as a negative control. **D.** Venn diagram showing the overlap of TET1 (blue) and NANOG (red) high confidence ChIP seq peaks in mouse ESCs. **E.** Gene ontology analysis performed on genes associated with TET1-NANOG co-bound sites. **F.** Genomic snapshots showing NANOG (green) and TET1 (red) ChIP-seq signals in the vicinity of NANOG transcriptional target genes.

